# Detecting Phenotypic Variability in *Lupinus angustifolius* Through Ecogeographic Land Characterization

**DOI:** 10.1101/2025.09.03.673914

**Authors:** C. Celdrán-Fernández, A. García-Fernández, F. J. Jiménez-López, M. L. Rubio-Teso, J.M. Iriondo-Alegría, N. González-Benítez, M.C. Molina-Cobos, C. Lara-Romero

## Abstract

Germplasm banks hold substantial potential across numerous branches of life sciences. However, this potential remains underutilized, partly due to the limited and incomplete characterization of the accessions. To address this issue, we propose the use of Ecogeographical Land Characterization (ELC) maps as a tool for classifying accessions into distinct ecological regions, thereby simplifying the characterization process. To test this approach, we collected *Lupinus angustifolius* seeds from multiple locations across the Iberian Peninsula, guided by a previously developed ELC map. Our results suggest that ELC maps effectively condense the Iberian landscape while capturing a moderate proportion of the species’ phenotypic variation. Overall, this approach offers a promising avenue to optimize germplasm conservation strategies while balancing resource limitations.

## Introduction

The number of accessions in germplasm banks, along with the rapid rate at which new accessions are continuously added, far exceeds the capacity for thorough characterization (FAO, 2025; Gonzalez et al., 2018; Yu et al., 2016). Without efficient characterization, the potential value of many accessions remains untapped, limiting their use. However, precise genotypic and phenotypic characterizations remain challenging. Identifying and measuring traits that are adaptive to specific environmental conditions is often labor and resource-intensive, requiring extensive fieldwork, laboratory and computational analyses (Soga & Gaston, 2025). Furthermore, the complex nature of genotype, phenotype, and environment interactions pose additional difficulties (Félix & Barkoulas, 2015; Gregorius, 1977). As a result, the full potential of germplasm collections remains largely underutilized, highlighting the need for innovative approaches to optimize and improve the characterization efforts, specifically in the current climatic context, where rapid and efficient responses are imperative.

Ecogeographical characterization emerges as an efficient tool to bridge this gap. This tool is based on the theorical relationship between the environmental characteristics of a site and features of plant populations occurring at that site, due to natural selection and local adaptation (Greene & Hart, 1999). The advancement of Geographic Information Systems, coupled with the growing availability of environmental data, has significantly enhanced the potential of this information as a predictive tool (Thormann et al., 2014). Various strategies have employed environmental data to facilitate the search for traits of interest. One notable example being the Focused Identification of Germplasm Strategy (FIGS) for plant breeding (Khazaei et al., 2013; Sunitha et al., 2024).

Among these, Ecogeographical Land Characterization (ELC) maps use bioclimatic, edaphic, and geophysical data, which is highly accessible, to model the adaptive scenarios in which a population has evolved (Parra-Quijano et al., 2011). ELC maps can be utilized to identify populations most likely to carry desired traits, thus, to simplifying the candidate selection process in plant breeding programs, particularly in the utilization of natural populations of crop wild relatives (CWR) (Garcia et al., 2017; González-Santos et al., 2024; Parra-Quijano et al., 2011; Rubio Teso et al., 2022; Thormann et al., 2014). Additionally, ELC maps are used in conservation biology, facilitating the identification of crucial areas or gaps germplasm collections for in *in-situ* or *ex-situ* conservation (Garcia et al., 2017; Ghamkhar et al., 2007; González-Santos et al., 2024; Parra-Quijano et al., 2012a, 2012b).

Ecogeographical modelling, including the use of ELC maps, is not without its limitations (Elith & Leathwick, 2009; Mateo et al., 2011). These models often assume linear and continuous species responses to environmental variables and are developed at specific spatial and temporal scales, with input data constrained by resolution or availability. Thus, these models can have limited predictive performance, particularly for biological traits (Anderegg, 2023; Lee-Yaw et al., 2022). While the relationship between phenotypic traits and ecogeographical data is fundamental to local adaptation, validation of ELC maps as predictors of phenotype remains scarce (Parra-Quijano et al., 2011; Steiner et al., 2001), thus limiting the broader applicability of this tool. Such validation is crucial for establishing the accuracy and reliability of ELC tools as suitable methods for classifying germplasm banks and supporting their applications.

This study employed *Lupinus angustifolius* L., commonly known as narrow-leafed or blue lupin, as a study species to evaluate the validity of ELC maps as a useful tool for predicting intraspecific variation. It examined phenotypic intraspecific variability across a large environmental range in the Iberian Peninsula, serving as a proof of concept for the tool’s ability to classify individuals based on climatic data from their environment of origin. *L. angustifolius* is a leguminous plant native to the Mediterranean region, commonly found in dry, sandy soils. This species holds not only ecological and evolutionary significance, but also agronomic value (Lemus-Conejo et al., 2023). Previous studies have shown phenotypic differences of this species in relation with geographical origin (Clements & Cowling, 1994) specially in the adaptation across ecosystems with variable aridity (Berger et al., 2020; Matesanz et al., 2022; Poyatos et al., 2023). The Iberian Peninsula hosts a gradient of environmental conditions (Giralt et al., 2017) and distinct *L. angustifolius* populations (Garg et al., 2022), making it an important area for exploring the species’ genetic diversity.

We hypothesized that the regions identified by the ELC map will be associated with differentiated climate adaptation strategies, determined by the climate of each ecological region. Therefore, a significant relationship is expected to be found between phenotypic traits and the climatic conditions defined by these. This approach will test whether ELC tools can capture local adaptations associated with climatic variation, providing a robust framework for modelling and predicting phenotypic traits from ecogeographical characterization of the territory.

## Materials and Methods

### Study species

*Lupinus* L., belonging to the Fabaceae family, contains 580 accepted species (POWO, 2024), making it one of the largest genera within this family. Among these, *Lupinus angustifolius* is a small, smooth-seeded Old World lupin that demonstrates a significant intraspecific variation (Wolko et al., 2011). It is predominantly self-pollinating, with a very low cross-pollination rate that typically falls between 0 and 3%, though this observation varies across studies (Dracup & Thomson, 2000; Williams, 1987). The lupin genome is characterized by high karyotypic variation, with chromosome numbers ranging from 2n = 32 to 52 (Wolko et al., 2011). For *L. angustifolius*, its genome is comprised of 20 pairs of chromosomes (2n = 40) (Garg et al., 2022).

### Ecogeographical Land Characterization (ELC) map generation

In this study, an ELC map of the Iberian Peninsula was constructed with the aim of identifying the ecological variables that best describe the distribution of *L. angustifolius*, as well as distinguishing differentiated regions that may harbor a broader variety of genotypic and phenotypic traits (Parra-Quijano et al., 2012a).

Ecogeographical variables used in this study were obtained from the Chelsa database and included 118 bioclimatic variables, along with latitude and longitude. Variable data was extracted from version 2.1, which has a 30 seconds (∼1 km2) spatial resolution (Karger et al., 2023). Distribution data was obtained from GBIF webpage (GBIF Backbone Taxonomy, 2023). Records of the same taxa found within a 1 km radius were removed, assuming they belonged to the same population (Rubio Teso et al., 2022).

The selection of variables was performed using a modified R script developed for the “SelecVar” tool of CAPFITOGEN3 (Parra Quijano et al., 2021). The R script gave each population of *L. angustifolius* a value for each bioclimatic variable, according to their geographic coordinates and assessed the importance of each variable in explaining the distribution of *L. angustifolius* populations. Variable importance estimates were calculated using a Random Forest Classification (RF) algorithm, with importance based on mean decrease accuracy (MDA) values (Cutler et al., 2007). The 15 bioclimatic variables with the highest MDA values were selected for further analysis. Following Garcia et al. (2017) and Rubio Teso et al. (2022), we retained the variables with the highest MDA value and removed all highly correlated variables (Pearson correlation coefficient > |0.5| and p < 0.05). Non correlated variables were used for the generation of the ELC map. The ELC map was generated following a modified R script of the ELC maps tool of CAPFITOGEN3. Map generation employed the clustering “elbow” method, which determines the cutoff points to identify the optimal number of clusters (Ketchen and Shook, 1996). The resulting map, with a 30 arc-sec resolution, was used to extract the ecogeographic regions (ELCr) using a modified R script of the “Representa” tool of CAPFITOGEN3 (Parra Quijano et al., 2021). Analyses regarding the generation of ELC maps were conducted using the R 3.6.3 version of the R environment (R Core Team, 2020) and scripts downloaded and adapted from CAPFITOGEN3 website.

### Seed collection

The previously constructed ELC map, which identified four ELCrs (see Results), was used to guide the sampling strategy. At least five collection sites were selected for each ELCr. At each site, 15-20 seeds were collected from a minimum of 20 individuals. All individuals in the same site were collected within a 1km radius, thus considering them the same population. To avoid choosing tightly related samples, a minimum distance of 3 meters was maintained between sampled plants. Seeds were collected between April and July of 2023, stored at room temperature until laboratory processing, after which they were kept at −20°C until sowing. The mean seed weight for each mother plant was determined by measuring ten seeds per mother plant. Collected seed material was deposited in Rey Juan Carlos University Germplasm Bank (https://ipt.gbif.es/resource.do?r=germoplasma-urjc).

### Set up of common garden experiment

A common garden experiment was conducted in Móstoles, Spain at the CULTIVE facilities of Rey Juan Carlos University (https://urjc-cultive.webnode.es/), to assess intraspecific trait variation. By unifying the environment, the experiment aimed to isolate and identify genetic differences among the regions (Kawecki & Ebert, 2004). The experimental design included 16 mother plants for each of the 20 collection sites, resulting in a total of 320 individuals. Sowing was carried out in November 2023 following the methods outlined by Sacristán-Bajo et al. (2023) and Poyatos et al. (2023). Briefly, for each mother plant, three seeds were sown after physical scarification in a 6L pot with a 1:1 mixture of commercial substrate and sand (Abonos Alonso S.A. Madrid, Spain). Germination and seedling emergence occurred in a greenhouse (25°-15°C and 70-45% humidity, with two waterings a day) before plants were transferred to outdoor conditions in January 2024, where they remained until senescence (June 2024). Watering was carried out on alternate days until April, and daily thereafter, aiming to maintain *ad libitum* conditions. In March 2024, once the seedlings were established, the biggest of the three seedlings per mother was selected to continue in the experiment.

### Trait measurement

#### Morphological traits

Seven morphological traits were measured: number of ramifications, leaflet biomass, root length, root collar diameter, number of secondary roots, root and shoot biomass, and root-to-shoot ratio. Biomass measurements were obtained after oven-drying the collected material. Leaflet biomass was calculated as the mean value of eight central leaflets. The root collar diameter was recorded at the point of separation between above- and below-ground biomass. The number of ramifications and secondary roots was determined by counting branches emerging from the main stem or main root. Except leaflet biomass, all morphological traits were measured at the end of their life cycle, leaflets were collected in April during vegetative growth.

#### Physiological traits

Relative Water content (RWC) and growth rate were measured as physiological traits. Leaflets collected for leaflet biomass were used to measure RWC following Jensen & Henson (1990) protocol:

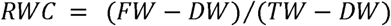

where FW, DW and TW refer to fresh, dry and turgid weight respectively.

Growth rate was calculated by calculating a regression coefficient based on four height measurements: during vegetative growth, pre-flowering, during flowering, and post-flowering at the end of the life cycle; following the general linear regression:

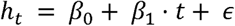

where *h_t_*is height in centimeters, *t* is time in days, β_0_ the intercept and β_1_ the growth rate.

#### Phenological and reproductive traits

The onset flowering was recorded as each plant reached the “*Diverging standard petal stage (anthesis)*” in accordance with the Protocrop manual for Lupin cultivation (Industry & Investment NSW, 2011). Reproductive traits included the number of pods, total number of seeds and seed weight. Seed weight was determined with all seeds weighted and divided by the total number of seeds.

### Data and statistical analysis

All statistical analyses were performed in R version 4.4.0 (R Core Team, 2024). Principal Component Analysis (PCA) was conducted to explore patterns of variation in phenotypic traits. The analysis was performed using the ‘prcomp’ function from ‘stats’ package in R (R Core Team, 2024). Prior to PCA, all continuous variables were standardized (mean-centered and scaled to unit variance) using the ‘scale’ function. To assess the effect of the grouping variable, PERMANOVA was conducted with 9999 permutations using the ‘vegan::adonis2’ function, based on Euclidean distance matrices calculated with the ‘vegan::vegdist’ function (Oksanen et al., 2001). Additionally, linear regression models were fitted for each PCA axis, using the climatic variable as the predictor, and the results were statistically evaluated through ANOVA analysis.

A linear mixed-effects model (LMM) was fitted for each phenotypic trait to evaluate the effect of the ELCr of origin. The general model structure was:

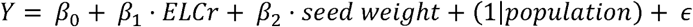

where Y represents the phenotypic trait, β_0_ is the intercept, β_1_ and β_2_ are the coefficients for each predictor variable (i.e., ELCr and seed weight), and E is the residual error term. The predictor variable (seed weight) is the mean seed weight of the mother plant, introduced to account for maternal effects. The random effect variable (population) refers to the site where seeds from the mother plant were retrieved from.

All traits were modelled using a Gaussian distribution, with the exception of count-based traits (number of secondary roots, number of branches, number of fruits, number of seeds), which were analyzed using generalized linear mixed models (GLMMs) with a Poisson distribution. Model fitting was performed using the ‘lme4’ package, with the ‘lmer’ function for continuous traits and the ‘glmer’ function for count data (Bates et al., 2015).

Outliers were identified based on Cook’s distance, calculated using the influence function from ‘stats’ package. Observations with Cook’s distance greater than 4/*n*, where *n* is the sample size, were considered influential and removed from the dataset. Model summaries were obtained using the ‘summary’ function, and Type II Wald chi-square tests were conducted using the ‘car::Anova’ function to assess the statistical significance of fixed effects. Pseudo R2 of the LMM was calculated with the same function, following Nakagawa & Schielzeth (2013) methodology. Post-hoc pairwise comparisons between levels of the variable ELC were performed using the ‘emmeans’ function from the R package ‘emmeans’ (Lenth, 2017), with the results adjusted using the False Discovery Rate (FDR) method.

Furthermore, a RF classification model was implemented to assess whether the phenotypic dataset was robust enough to train a predictive model capable of classifying individuals based on their ecogeographical region of origin. The analysis was conducted using the ‘randomForest’ package in R (Liaw & Wiener, 2002). Phenotypic traits were standardized, and the dataset was split into training (80%) and testing (20%) sets using stratified sampling to preserve the distribution of ELC categories. The number of trees in the random forest was selected using the Out of Bag score. Thus, the algorithm was trained with 1000 trees. The Random Forest model was trained 999 times using a different randomization of the data, and functional metrics were calculated as the average of all the iterations. Key classification metrics were calculated, including accuracy, precision, recall and specificity. Importance of each phenotypic trait in the classification task was assessed using the MDA metric. Traits with a higher MDA are considered more influential.

## Results

During the ELC map construction, bioclimatic variables were ranked by their MDA and the top fifteen were selected. These variables were highly correlated (Table S1). Removal of correlated variables resulted in one selected variable, June’s Climatic Moisture Index (CMI). CMI measures the balance between the amount of water added to the environment through precipitation and the amount of water lost through evapotranspiration. The generation of the ELC map for the Iberian Peninsula using this variable resulted in a map with four ELC categories (Fig. 1). These ecological regions (ELCr) were denoted from 1 to 4, corresponding to increasing values of June’s CMI. The resulting ELC map guided the selection of twenty sites across ELCr 1 to 3 (see Fig. 1). Not enough samples from ELCr 4 could be retrieved, consequently, it was excluded from further analysis.

**Figure 1.**
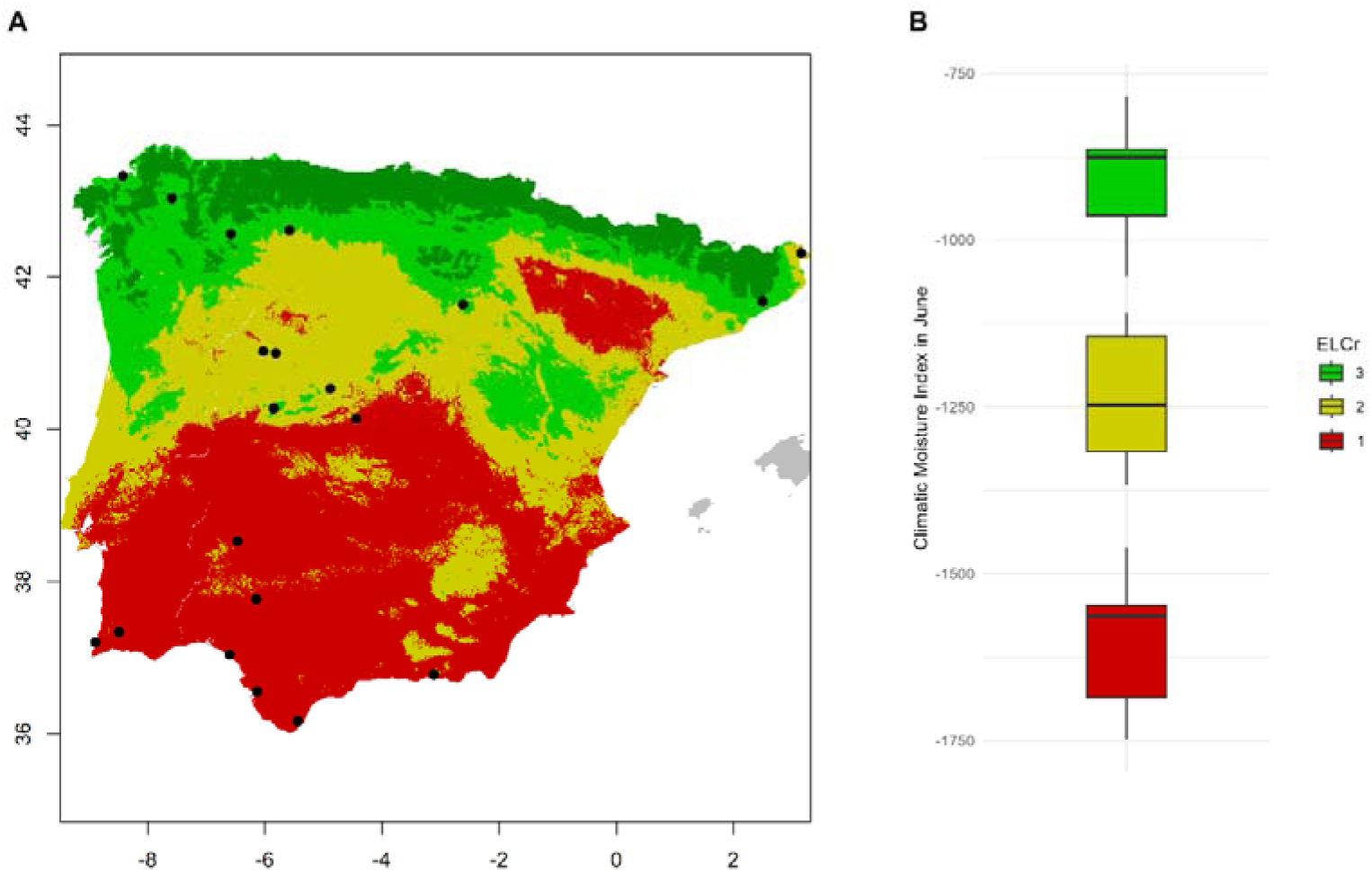
Ecogeographic Land Characterization Map of *Lupinus angustifolius*. (A) ELC map was generated using June’s Climatic Moisture Index (CMI). Four distinct ELCrs were defined by the elbow algorithm. Collection sites for the common garden are marked with filled dots. (B) Box plot showing the distribution of June’s CMI values across collection sites.

In total, 14 phenotypic traits were measured. PCA revealed that the first three components explained 29.33%, 15.61%, and 10.08% of the total variation, respectively. The primary contributors to PC1 were morphological traits, including root and shoot biomass and growth rate. PC2 was influenced by both morphological and reproductive traits, such as the number of fruits and seeds, root-to-shoot ratio, and number of branches. In contrast, PC3 was predominantly shaped by reproductive traits like flowering onset and seed biomass, as well as root length (Fig. S3). The linear model ANOVA identified significant correlations for PC1 (p < 0.0001=, r² = 0.1) and PC3 (p = 0.022, r² = 0.03). Similarly, the PERMANOVA analysis confirmed significant associations for PC1 (p < 0.0001, r² = 0.12) and PC3 (p = 0.019, r² = 0.05) (Fig. 2).

**Figure 2.**
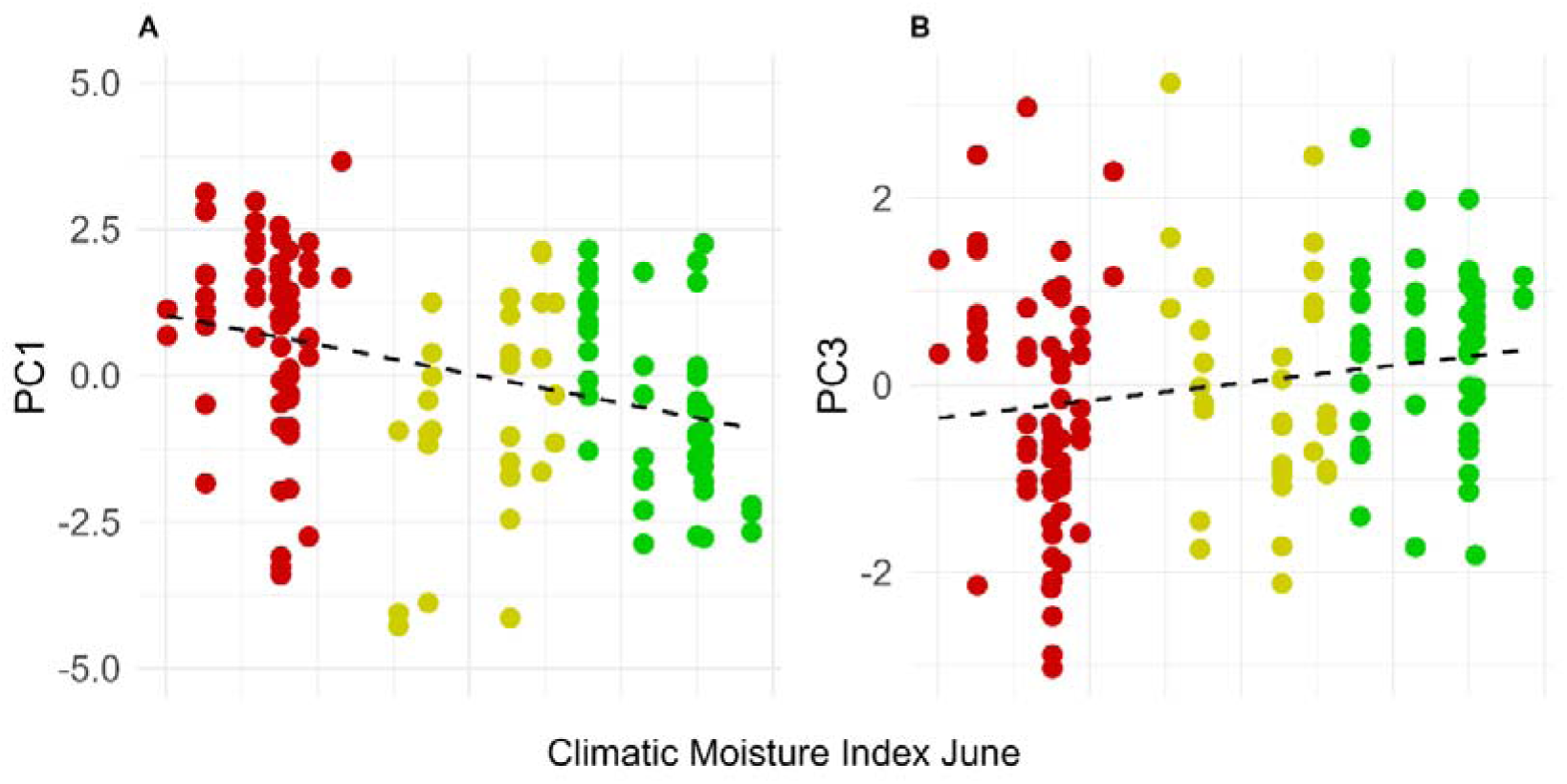
Principal Component Analysis of *Lupinus angustifolius* traits and their relationship with June’s CMI. (A) The first principal component (PC1) is mainly associated with morphological traits (root and shoot biomass and growth rate), while (B) the second principal component (PC3) is primarily influenced reproductive traits (flowering and seed biomass).

The LMMs and GLMs used to investigate the effect of ELCr on the intraspecific variation of the measured traits identified eight traits as significant: leaflet biomass (p = 0.003, r² = 0.24), root length (p = 0.002, r² = 0.11), root collar diameter (p = 0.01, r² = 0.26), number of secondary roots (p < 0.001, r² = 0.34), number of fruits (p = 0.04, r² = 0.10), shoot biomass (p < 0.00001, r² = 0.35), root biomass (p < 0.0001, r² = 0.31), and growth rate (p = 0.01, r² = 0.27) (Table S2). Post-hoc analysis revealed that ELCr 1 showed notable differences compared to ELCr 3 for all traits identified in the mixed models, except for the number of fruits. Similarly, ELCr 1 differed from ELCr 2 for all these traits, except for root length, number of fruits, and growth rate (Fig. 3). No clear trend in trait expression was observed between ELCr 2 and ELCr 3, except for the number of fruits, which varied between these two groups.

**Figure 3.**
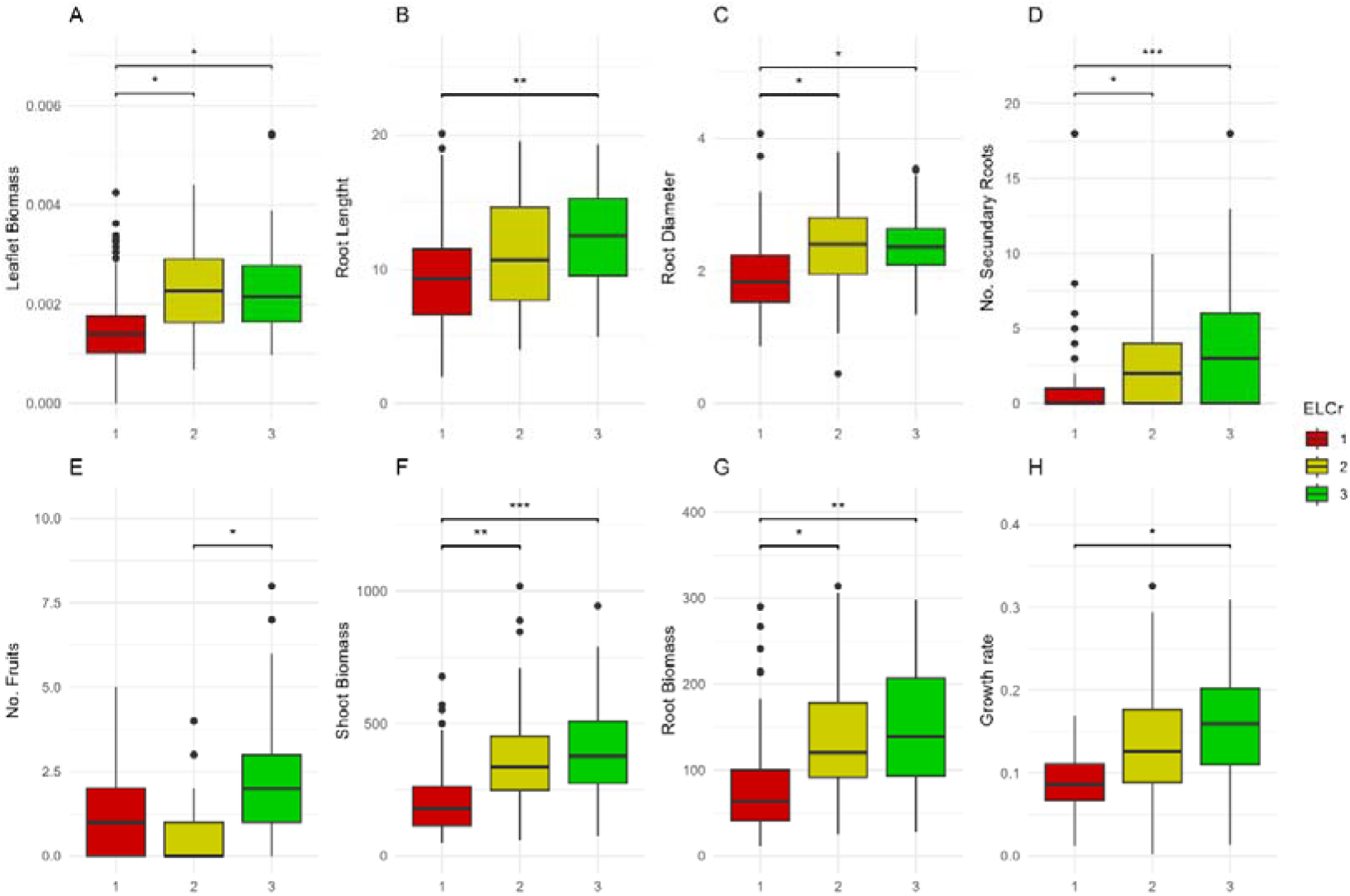
Boxplots of significant traits across Ecogeographic Land Characterization regions (ELCr). Eight traits were identified as significantly influenced by ELCr in the mixed models: leaf biomass, root length, root collar diameter, number of secondary roots, number of fruits, shoot biomass, root biomass, and growth rate. . Significant differences between ELCr groups are indicated by asterisks based on post-hoc tests: p < 0.05, p < 0.01, p < 0.001.

The RF trained to classify individuals based on their phenotypic data achieved an overall average accuracy of 73.35%, with 54.16% precision, 55.76% recall and 79.90% specificity. Regarding individual class metrics, the algorithm showed notably lower precision and recall for classifying ELCr 2, with values of 21.34% and 43.11%, respectively. The remaining metrics for individual classifications ranged from 60 and 80% (Table S3). Additionally, ELCr 2 was underpredicted compared to the other ELCrs (Fig. S3). This underrepresentation appears to be compensated by an overprediction of ELCr 1, which tends to absorb individuals that are likely to belong to ELCr 2.

Random Forest overall importances (Fig. 4) showed leaflet biomass as the trait with a highest importance (18.92%). Followed by growth rate, flowering onset, number of secondary roots, seed weight and shoot biomass with a contribution to MDA greater than 10%, which aligns with the traits identified as significant in the linear mixed model analysis. Four traits had an MDA lower than 5%: root length, root-to-shoot ratio, number of branches and RWC. These traits, except for root length, also coincide with those that showed the highest p-values in the linear mixed model. Individual class importances, measured as the MDA for the classification of each class label (ELCr, in this case), were lower for most traits when classifying ELCr 2, with a negative MDA observed for five of the traits. For seven traits, ELCr 1 had the highest class-specific MDA, particularly for those traits that held greater importance. ELCr 3 class-specific importances had closer values to those of ELCr 1, with higher values for less important traits (Fig. 4).

**Figure 4.**
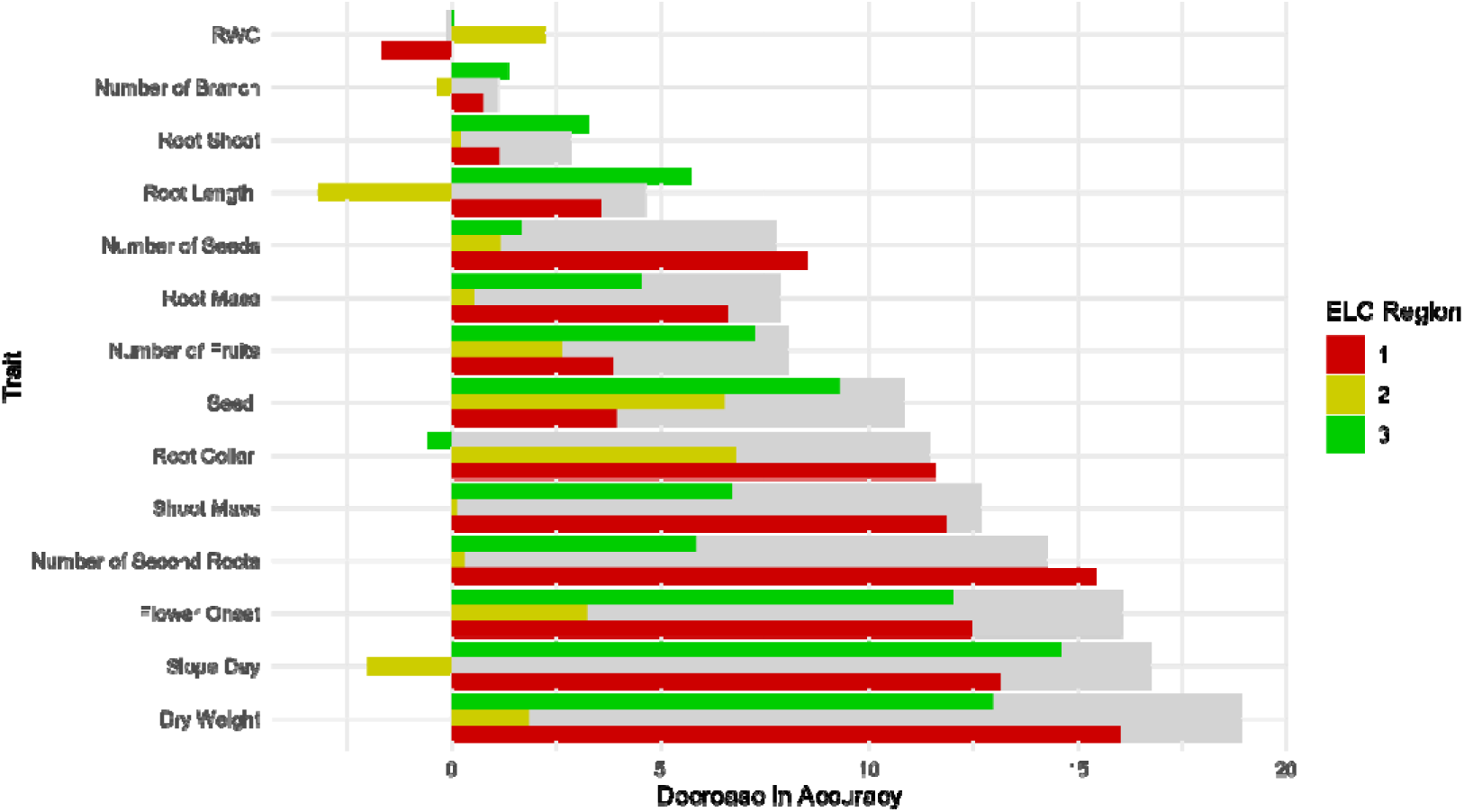
Importance of traits in the Random Forest classification model. Random Forest model for classification of individuals into the three ELCr based on phenotypic traits. The grey bars represent the Mean Decrease in Accuracy for each trait, while colored bars indicate the Mean Decrease in Accuracy for each ELC category and trait.

## Discussion

Our study aimed to assess whether the regions identified by the ELC map, based on climatic data, correspond to differentiated phenotypes in *Lupinus angustifolius* populations. We hypothesized that populations from distinct ELCr would exhibit significant phenotypic differences linked to their characteristic climates, supporting the use of ELC as a predictive tool for local adaptation. ELC map constructed with June’s CMI, delimited ELCr capturing significant phenotypic variation between populations.

### Iberian *Lupinus angustifolius* intraspecific variation

Phenotypical analysis revealed significant differences among *L. angustifolius* retrieved from different ELCr across the Iberian Peninsula, suggesting population differentiation and evolutionary diversification. Since all individuals grew on a common garden phenotypic differentiation is the result of genetic differentiation and maternal effects. Maternal effects were accounted for in the LMM by including mother plant’s average seed weight in it.

Phenotypic observations align with previous descriptions of drought adaptation in the genus *Lupinus*. Populations in drier areas typically show a more conservative strategy and, therefore, individuals show lower values for morphological traits. This aligns with our findings from the LMM analysis and RF trait importance results. However, other expected adaptations such as earlier flowering time or fewer but larger seeds (Berger et al., 2020; Poyatos et al., 2023; Talhinhas et al., 2006), did not reach statistical significance in the LMM-ANOVA analysis. In spite of this, both traits played an important role in the RF algorithm (MDA > 10%), indicating that, while they lacked statistical significance, they exhibit enough meaningful variation for the algorithm to differentiate between groups. This discrepancy can be attributed to the different approaches each analysis takes toward the variables. ANOVA-based analyses treat each variable independently, detecting differences in mean values between groups to test the null hypothesis (Kim, 2014). In contrast, RF algorithm creates a predictive model accounting for variable interactions and non-linear relationships (Grömping, 2009; Louppe, 2014). As a result, some variables may exhibit high MDA values even without significant differences between groups. Conversely, statistically significant traits can show low MDA values due to having their predictive capability absorbed by another variable.

Some non-significant and low MDA traits also provide insight into adaptation patterns. The results obtained for root-to-shoot ratio suggest that there is no clear adaptation towards larger root systems in drier climates. This could indicate that *L. angustifolius* does not rely on an extensive root system as a primary drought adaptation strategy, or that selection for this trait is weak across the studied populations. However, Berger et al. (2020) reported significant differences in root-to-shoot ratios between high- and low-rainfall areas in Mediterranean genotypes. These differences may stem from the methods used to calculate the ratio. While we measured biomass at the end of the plant’s life cycle, when leaves had already senesced, Berger et al. (2020) harvested material during pod set, allowing for leaf biomass to be included. Another factor that may contribute to this discrepancy could be differences in how categories were built: high- and low-rainfall, and ELCrs in our case. Moreover, RWC also did not show significant differences among ELCr. However, this result is not necessarily unexpected. When plants are well-watered, RWC is typically maintained within a relatively narrow range due to homeostatic mechanisms. Significant differences in RWC may only become evident under drought stress, where phenotypic plasticity plays a stronger role in adaptation (Berger et al., 2020; Jensen & Henson, 1990; Pinheiro et al., 2004).

### ELC as predictive tools for germplasm banks, resource conservation, breeding programs and other applications

PCA confirms that overall phenotypic data correlates with June’s CMI, showing it is, as a single variable, a useful predictor for intraspecific variation in Iberian *L. angustifolius*. In total, 39.4% of the variation has a significant relationship with this climatic variable. Key traits in these components include root and shoot biomass, growth rate, flowering onset and seed mass, which, as discussed previously, show an important relationship with the region of origin. In contrast, PC2 did not show a significant relationship with June’s CMI, which is consistent with the non-significant associations observed for the traits driving this component, number of seeds, root-to-shoot ratio and number of branches.

In the context of site selection and germplasm collection, it is, however, important to establish clearer territorial limits. Thus, we developed a climatic variable-based regional classification identifying four distinct regions, which is a significant simplification of the Iberian Peninsula complex landscape. Phenotypic variation among groups showed most of the traits being significantly influenced by ELCr origin and was able to explain a moderate proportion of trait variation —based on marginal pseudo-R² values— supporting the classification’s relevance.

Training a random forest classifier on the phenotypic data tests whether phenotypic differences are sufficient for reliable classification into the three ELCr, rather than resulting in a random assignment. The model obtained moderate to high accuracy, suggesting phenotypic differences between groups substantial enough to be captured by a RF algorithm. However, the model’s lower precision and recall, particularly for ELCr 2, indicate that not all groups are equally well defined. In contrast, ELCr 1 and 3 were more consistently identified, pointing to clearer phenotypic boundaries.

Interestingly, ELCr 2 consistently shows lower importance for most influential traits and its underpredicted in the RF model. Given the limited size of our dataset, and the intermediate nature of ELCr 2, the RF model likely struggled to identify influential interactions specific to this group. Notably, while the LMM post-hoc analyses revealed greater confusion between ELCr 2 and 3, the RF model primarily misclassified individuals between ELCr 1 and 2. This phenomenon is consistent with ecological expectations: intermediate regions often show mixed or intermediate responses due to mixed adaptive pressures, gene flow (Sacristán-Bajo et al., 2025) or plasticity (Matesanz et al., 2022). Such patterns indicate that the discriminative power of the ELC maps may be reduced when ELCr are climatically similar, as comparable selection pressures naturally constrain phenotypic differentiation. Yet, when regions are more distinct, the ELC framework remains effective in capturing clearer phenotypic divergence. Additionally, it is possible that genotypic or demographic differences exist within ELCr 2 that were not captured in this analysis. Despite this, the approach effectively captures phenotypic differentiation with a very simple classification: three groups based in one climatic variable.

ELC maps and other ecogeographical modelling approaches, such as FIGS (Khazaei et al., 2013) and predictive characterization techniques (Thormann et al., 2014), have focused on targeted characterization of germplasm banks accessions, meaning they identify specific traits of interest rather than providing a general classification of the collection (Sunitha et al., 2024). In contrast, our approach proposes a simplified version of an ELC map, organizing the territory into distinct regions—ELCrs—allowing all individuals of the studied species to be classified into one of these regions. Consequently, this technique enables a unified classification of all accessions of a given species, in a given territory, across different germplasm banks. As previously discussed by Anderegg (2023) and Lee-Yaw et al. (2022), ecogeographical modelling has limitations in predicting biological differentiation, thus cannot replace large-scale genomic and phenotypic assessments, but it offers a practical starting point to an ongoing need.

## Acknowledgements

The authors declare no conflicts of interest regarding this manuscript. The authors acknowledge the support from the Writing retreats IICG-URJC 2025 call.

