## Supplementary Table 1 for "Detecting Phenotypic Variability in *Lupinus angustifolius* Through Ecogeographic Land Characterization"

**Supplementary Table 1.** **ANOVA results from Linear Mixed Models (LMM) examining the effect of Ecogeographic Land Characterization regions (ELCr) on traits.** The table includes the Pseudo R² (marginal), Chi-square statistic, p-value for the ELC effect, and significance levels. Asterisks indicate significance: p ≤ 0.05 (*), p ≤ 0.01 (**), and p ≤ 0.001 (***). Traits with non-significant p-values are marked as "ns".

| Trait | Marginal R2 | Chi Square | p-value | Significance |
| --- | --- | --- | --- | --- |
| Onset flowering | 0,112 | 3,307 | 1,91e-01 | ns |
| RWC | 0,009 | 1,352 | 5,09e-01 | ns |
| Leaflet biomass | 0,236 | 11,311 | 3,50e-03 | ** |
| Root length | 0,109 | 12,472 | 1,96e-03 | ** |
| Root collar diameter | 0,258 | 9,156 | 1,03e-02 | * |
| No. Secondary roots | 0,338 | 14,641 | 6,62e-04 | *** |
| No. Branch | 0,007 | 0,159 | 9,24e-01 | ns |
| No. Fruits | 0,102 | 6,031 | 4,90e-02 | * |
| No. Seeds | 0,070 | 1,905 | 3,86e-01 | ns |
| Seed weight | 0,218 | 1,381 | 5,01e-01 | ns |
| Shoot biomass | 0,354 | 24,690 | 4,35e-06 | *** |
| Root biomass | 0,308 | 18,660 | 8,87e-05 | *** |
| Root-shoot ratio | 0,001 | 0,016 | 9,92e-01 | ns |
| Growth rate | 0,302 | 10,844 | 4,42e-03 | ** |
