## Supplementary Table 2 for "Detecting Phenotypic Variability in *Lupinus angustifolius* Through Ecogeographic Land Characterization"

**Supplementary Table 2. Random Forest Metrics. The table includes** the metrics of the overall model, referred as mean, and for the individual classes. It includes Accuracy, precision, recall and specificity.

| Class | Accuracy | Precision | Recall | Specificity |
| --- | --- | --- | --- | --- |
| Mean | 73,35 | 54,16 | 55,76 | 79,90 |
| ELCr 1 | 69,44 | 77,54 | 64,33 | 76,82 |
| ELCr 2 | 76,15 | 21,34 | 43,11 | 80,35 |
| ELCr 3 | 74,46 | 63,60 | 59,84 | 82,52 |
