## Supplementary figures and images for "Detecting Phenotypic Variability in *Lupinus angustifolius* Through Ecogeographic Land Characterization"

### Supplementary Figure 1

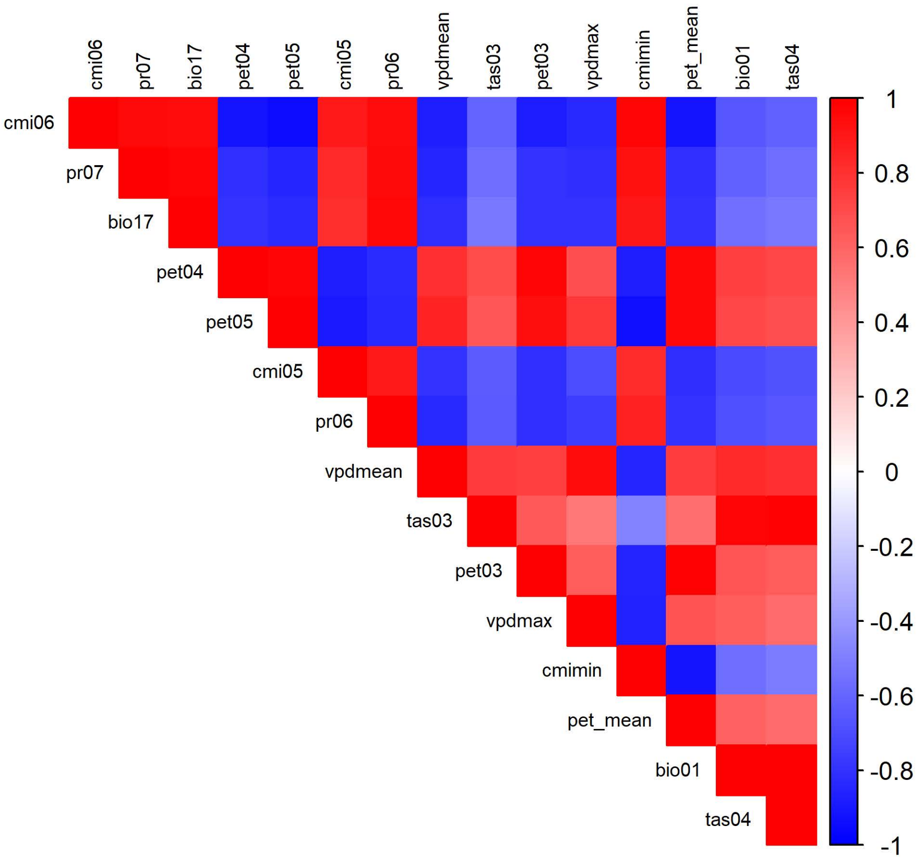

### Supplementary Figure 2

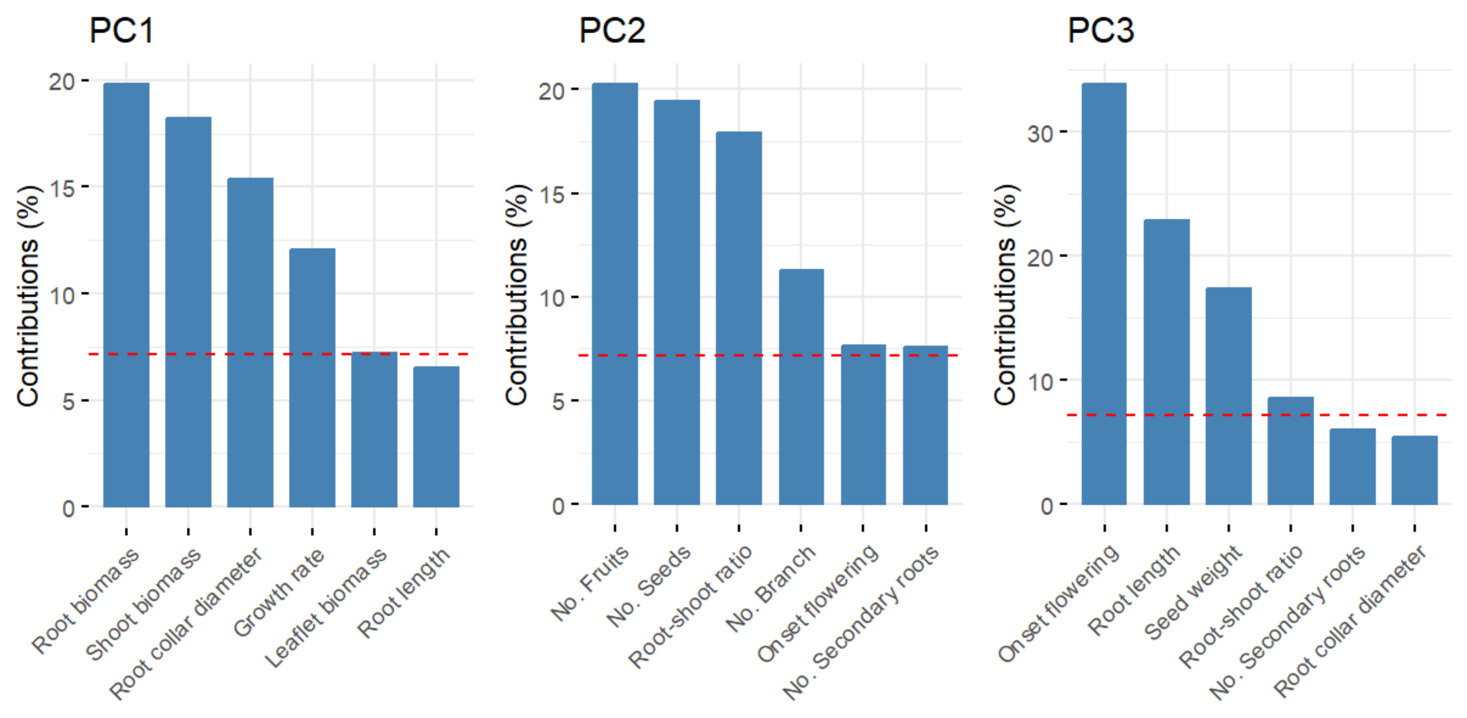

### Supplementary Figure 3

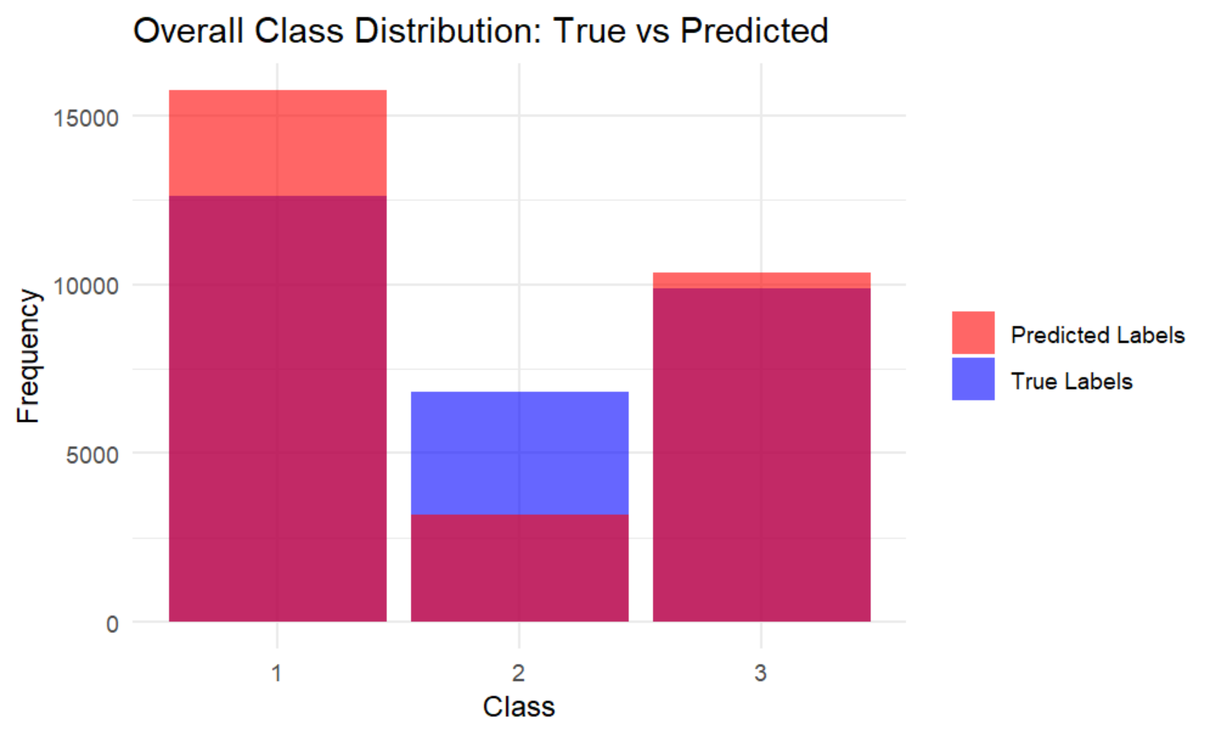
